# Netrin-1 regulates the balance of glutamatergic connectivity in the adult ventral tegmental area

**DOI:** 10.1101/2022.05.09.491091

**Authors:** Marcella M Cline, Barbara Juarez, Avery C. Hunker, Ernesto G. Regiarto, Bryan Hariadi, Marta E. Soden, Larry S. Zweifel

## Abstract

The axonal guidance cue netrin-1 serves a critical role in neural circuit development by promoting growth cone motility, axonal branching, and synaptogenesis. Within the adult mouse brain, expression of the gene encoding netrin-1 (*Ntn1)* is highly enriched in the ventral midbrain where it is expressed in both GABAergic and dopaminergic neurons, but its function in these cell types in the adult system remains largely unknown. To address this, we performed viral-mediated, cell-type specific CRISPR-Cas9 mutagenesis of *Ntn1* in the ventral tegmental area (VTA) of adult mice. *Ntn1* loss-of-function in either cell type resulted in a significant reduction in excitatory postsynaptic connectivity. In dopamine neurons, reduced excitatory tone had a minimal phenotypic behavioral outcome; however, reduced glutamatergic tone on VTA GABA neurons induced behaviors associated with a hyperdopaminergic phenotype. Loss of *Ntn1* function in both cell types simultaneously largely rescued the phenotype observed in the GABA-only mutagenesis. These findings demonstrate an important role for netrin-1 in maintaining excitatory connectivity in the adult midbrain and that a balance in this connectivity within two of the major cell types of the VTA is critical for the proper functioning of the mesolimbic system.

## INTRODUCTION

Proper regulation of the midbrain dopamine system is essential for numerous brain functions and behavior^1^. Disruption in the balance of midbrain dopamine neuron activity has been linked to several neurological and psychiatric conditions, including autism^2^, schizophrenia^3^, and substance use disorders^4^. Within the VTA, the activity of dopamine neurons is regulated in part by inhibitory (GABAergic) and excitatory (glutamatergic) synaptic input. The molecular mechanisms that maintain inhibitory and excitatory connectivity in the adult midbrain, however, remain poorly resolved.

Genome-wide association studies and analysis of *de novo* mutations has strongly implicated genes regulating neuronal axon guidance in neurodevelopmental disorders^5,6^. Although the impact of mutations in these genes early in development is likely critical for their role in neurodevelopmental disorders, many of the genes maintain high levels of expression in the adult brain and their functions in this context is less understood. We previously demonstrated that the axonal guidance receptor Robo2 is necessary for the maintenance of inhibitory synaptic connectivity in the adult VTA^7^, suggesting that axonal guidance proteins have a critical function in maintaining synaptic connectivity in the adult midbrain.

Netrin-1 is predominately recognized for its role in neurodevelopmental processes^7-11^. During development the gene encoding netrin-1 (*Ntn1*) is highly expressed throughout the central nervous system (CNS). Following this critical period global expression decreases^8^, but expression within the limbic system, particularly in the ventral midbrain persists. Consistent with the continued function of netrin-1 following early development, genetic inactivation of either *Ntn1*^9^ or its receptor *Dcc*^10^ from forebrain glutamatergic neurons in late postnatal development results in significantly impaired spatial memory in adult mice that corresponds to a loss of hippocampal plasticity. Within the VTA, *Dcc* expression levels in adult mice are significantly upregulated following amphetamine exposure^11^, and *Dcc* haploinsufficient mice display blunted locomotor response to amphetamine^12^, consistent with increased excitatory synaptic strength in the VTA following amphetamine treatment^13^. These results suggest a potential role for netrin-1 signaling through Dcc in regulating excitatory tone in the adult dopamine system.

To determine whether netrin-1 regulates excitatory synaptic connectivity in the VTA of adult mice, we used viral-mediated, Cre-inducible CRISPR/Cas9^14^ to selectively mutate *Ntn1* in midbrain dopamine and GABA neurons. We find that *Ntn1* loss of function significantly reduces postsynaptic glutamate receptor-mediated currents in a cell-autonomous manner similar to what has been reported previously in the adult hippocampus^10^. We further show that *Ntn1* loss of function in VTA GABA neurons has a more profound effect on behavior than loss of function in VTA dopamine neurons. Intriguingly, simultaneous loss of function of *Ntn1* in both cell types of the VTA largely rescues the behavioral phenotypes observed following mutagenesis in VTA GABA neurons alone. Collectively, these data demonstrate that the balance of excitatory synaptic connectivity onto VTA dopamine and GABA neurons is critical to the function of the mesolimbic system and that netrin-1 plays an important role in this process.

## RESULTS

### *Ntn1* expression and mutagenesis in the VTA

*In situ* hybridization analysis of *Ntn1* from the Allen Institute mouse brain expression atlas^15^ shows diffuse and low levels of expression throughout the adult mouse brain, with moderate expression levels in the cerebellum and hippocampus (Figure 1A), and the highest level of expression in the ventral midbrain (substantia nigra and ventral tegmental area). The VTA is comprised of multiple cell types ^16^; to determine the cell type specific expression of *Ntn1* within the heterogeneous VTA, we performed RNAscope *in situ* hybridization on midbrain slices from adult wild-type mice (>8 weeks of age) and probed for *Ntn1, Th* (tyrosine hydroxylase, a marker of dopamine neurons) and *Slc32a1* (vesicular GABA transporter [Vgat], a marker of GABA neurons). We found *Ntn1* expression to be present throughout the VTA, largely localized to *Th*-positive neurons but also present in GABA neurons (Figure 1C-F). *Ntn1* expression co-localized with *Th* expression in dopamine producing neurons (64% co-localization) and *Slc32a1*-expressing GABA neurons (30% co-localization) (Figure 1F). The remaining 6% of *Ntn1* expressing cells that do not co-localize with *Th* or *Slc32a1* are likely glutamatergic neurons^16^. Immunohistochemistry for netrin-1 and Th (Figure 1G) confirmed the presence of netrin-1 in dopamine and non-dopamine producing (Th-negative) cells.

**Figure 1.**
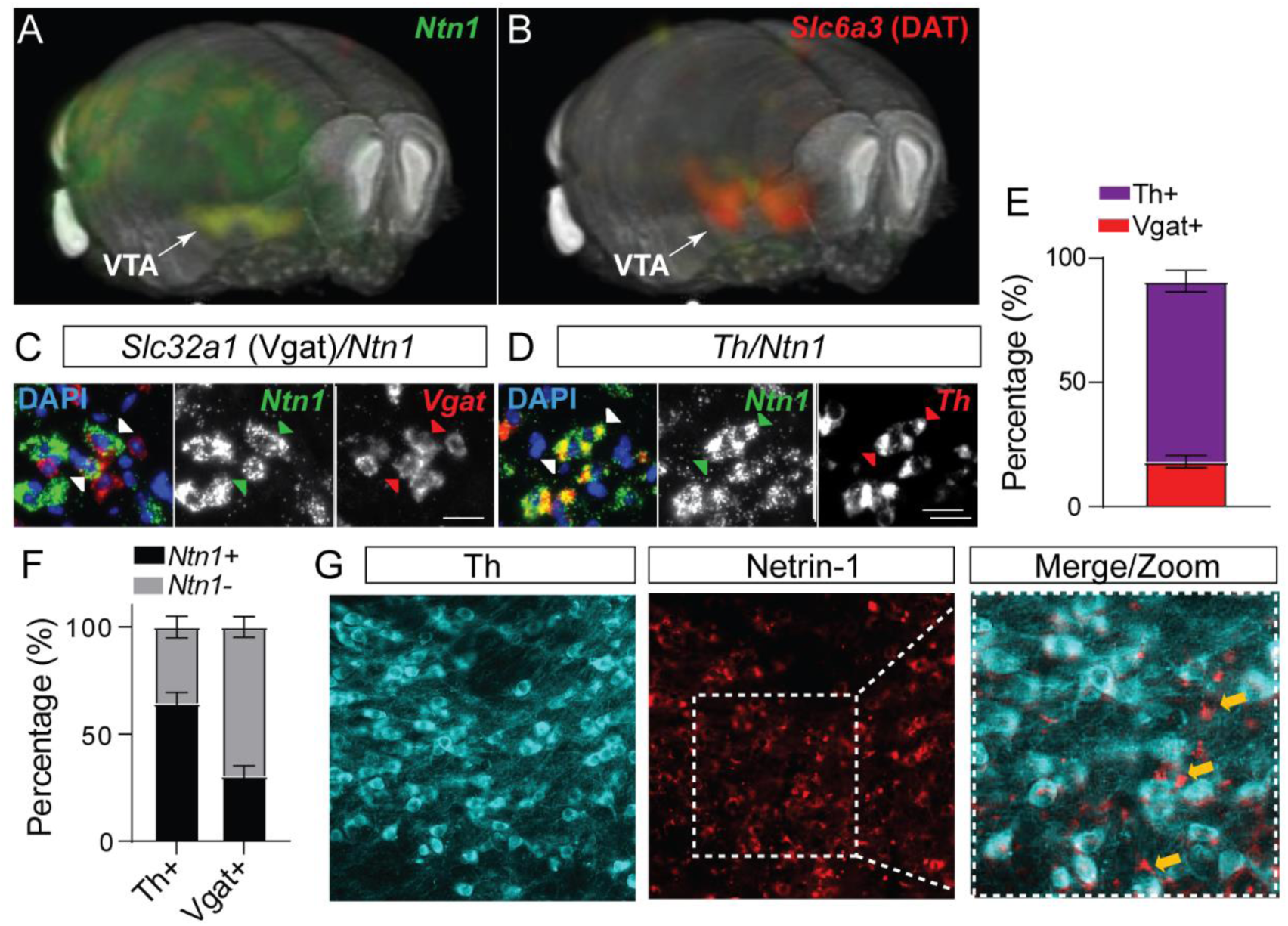
Netrin-1 is present in the adult VTA, and expressed by both dopamine and GABA neurons. 3D display of *Ntn1* (A) and *Slc6a3* (B, dopamine marker) from the Allen Brain Atlas. (C-D) 20X magnification images of in situ hybridization (RNAScope) for *Ntn1* (green) and *Slc32a1* (GABA marker; red, C) and *Th* (dopamine marker; red; D). Arrows indicate co-labeling of *Ntn1* with *Slc32a1* (C) or *Th* (D). Scale bar indicates 20μm. (E-F) Quantification of cell type expression. Of the cells expressing *Ntn1*, 72.2% were dopaminergic (*Th+*) and 18.1% were GABAergic (*Slc32a1+;* (E). (F) Of the total of *Th+* identified cells, 64.5% co-expressed Ntn1 (35.6% did not expressed Ntn1), and 30.4% of *Slc32a1* identified cells co-expressed *Ntn1* (69.5% did not expressed Ntn1). (G) Immunohistochemistry confirms the presence of Netrin-1 protein (red) in both Th+ (cyan) and non-dopamine cells (Th-cells, indicated by yellow arrows).

To selectively mutate *Ntn1* in specific cell types in the VTA, we designed a single guide RNA (sgRNA) targeting exon 2 in mice (sg*Ntn1*; Figure 2A) and cloned it into an AAV packaging plasmid containing a Cre-recombinase dependent expression cassette for SaCas9^14^. To determine the efficiency of *Ntn1* mutagenesis, we injected DAT-Cre (DAT^IRES^Cre; *Slc6a3*^Cre/+^) mice (aged 8-10 weeks) bilaterally into the VTA with either AAV-FLEX-SaCas9-HA-sg*Ntn*1 and AAV-FLEX-YFP (DAT-Cre *Ntn1*-cKO mice) or AAV-FLEX-SaCas9-sg*Rosa26* (a gene locus with no known function; control mice). Four to five weeks following injection, we performed immunohistochemistry for netrin-1 and Th. *Ntn1* conditional knockout (cKO) resulted in a significant reduction (∼80%) in the proportion of VTA Th-positive cells co-labeled with netrin-1 in DAT-Cre *Ntn1* cKO mice compared to controls (Figure 2D-E).

**Figure 2.**
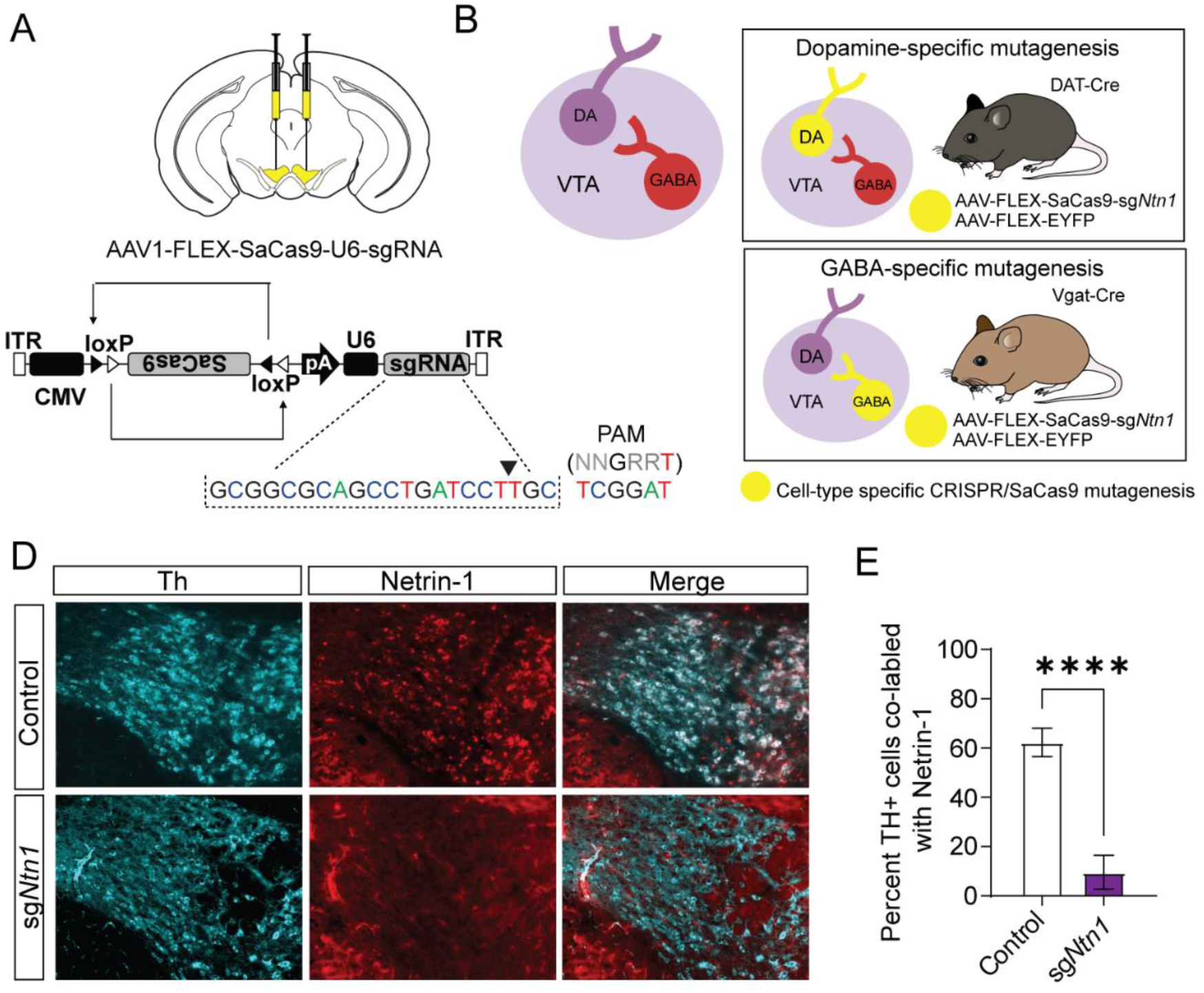
Virally delivered CRISPR-Cas9 complex targeting the *Ntn1* locus results in significant reduction in Netrin-1 antibody staining. (A-B) Schematics summarizing cell type specific knockout procedure. (A) Adult mice were injected bilaterally into the VTA with AAV-FLEX-SaCas9-HA-sg*Ntn1* and AAV-FLEX-YFP. Control mice received an equivalent volume of -sgRosa26 and/or AAV-FLEX-YFP. SaCas9 is virally delivered into the genome in the inactive orientation and returned to the active orientation only in the presence of Cre recombinase, limiting Cas9 expression to target cells. (B) Schematic of the VTA (left) showing VTA GABA neurons project to and inhibit VTA dopamine neurons. By using transgenic Cre-driver mouse lines (right) viral delivery of SaCas9 results in gene disruption in specifically VTA dopamine neurons (DAT-Cre mice, top panel), or VTA GABA neurons (Vgat-Cre mice, bottom panel). (D) Example images for Th (cyan) and Netrin-1 (red) immuno staining in the VTA of mice injected with control or sgNtn1 CRISPR virus. (E) Quantification of the percentage of Th+ cells co-labled with Netrin-1 (t=8.179, df=10, 62.25 ±5.796 vs 9.586±2.807, ****p<0.0001).

### Netrin-1 regulates excitatory connectivity within the adult VTA

Previous research has shown netrin-1 regulates excitatory synaptic connectivity in the adult hippocampus^17^. To determine the impact of *Ntn1* loss of function on synaptic connectivity, DAT-Cre or Vgat-Cre mice were injected with AAV1-FLEX-SaCas9-U6-sg*Ntn1* and AAV1-FLEX–YFP (Figure 3A and 3E). After at least 4 weeks, miniature excitatory postsynaptic currents (mEPSCs) were recorded from fluorescently identified dopamine or GABA neurons of the VTA. *Ntn1* mutagenesis in dopamine neurons resulted in significantly reduced mEPSC amplitude and frequency (Figure 3B-D). Similarly, *Ntn1* mutagenesis in VTA GABA neurons also resulted in significantly reduced mEPSC amplitude and frequency (Figure 3F-H). We did not detect significant effects on miniature inhibitory postsynaptic currents (mIPSCs) in VTA dopamine or GABA neurons following *Ntn1* mutagenesis in these cells (Figure 3 supplement 1), suggesting netrin-1 does not play a role in regulating inhibitory connectivity in these cells.

**Figure 3.**
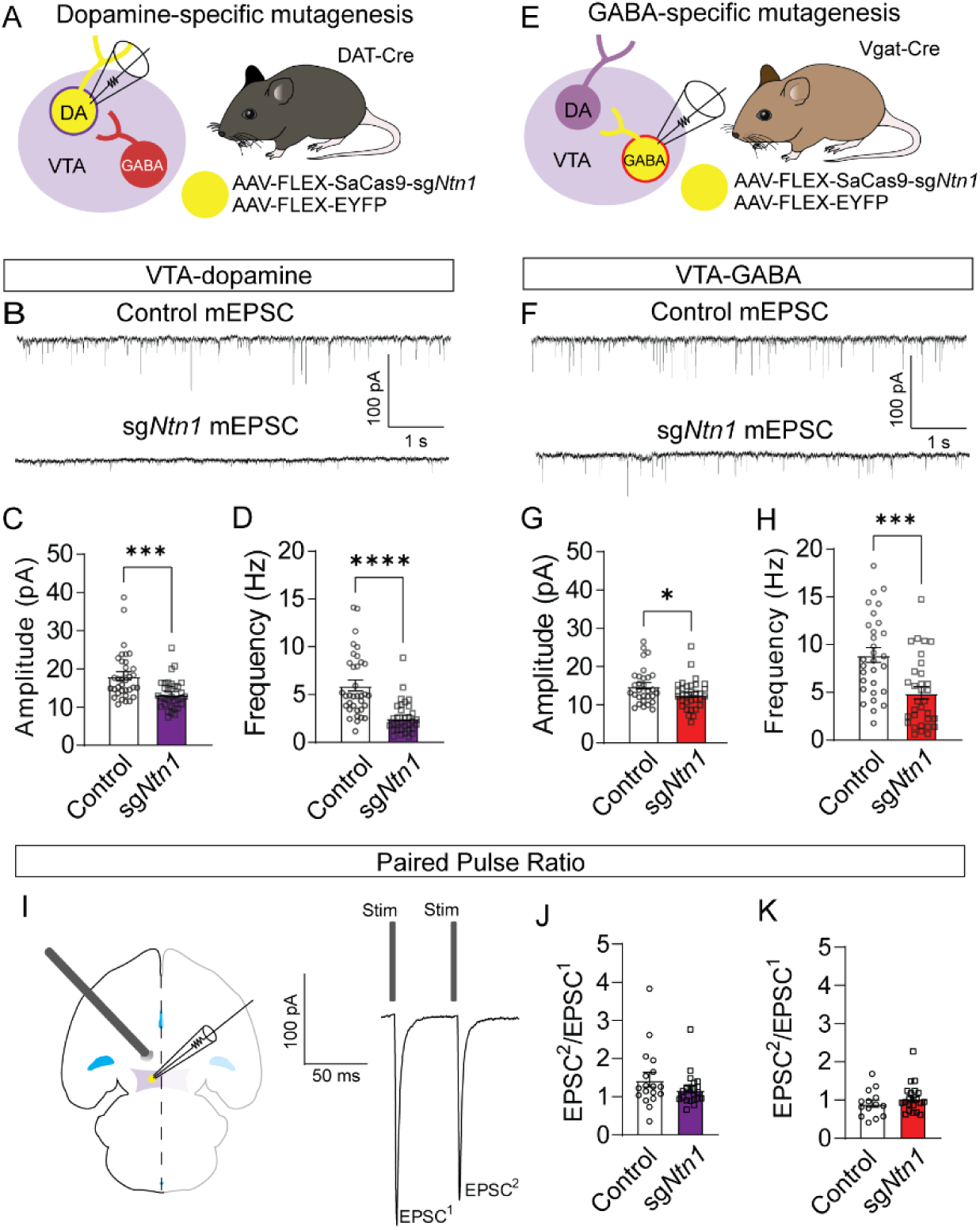
Loss of *Ntn1* results in significant reduction in excitatory post synaptic current. (A) Schematic of DAT-Cre dopamine specific Ntn1cKO. (B) Sample traces from control (top panel) and DAT Ntn1 cKO mice (bottom panel). (C-D) mEPSC amplitude (C) and frequency (D) measured from fluorescently identified dopamine neurons (n=35 controls, n=33 cKO, t=3.744, df=66, ***p<0.001 and t=5.259, df=66, ****p<0.0001). (E) Schematic of Vgat-Cre GABA specific Ntn1cKO. (F) Sample traces from control (top panel) and Vgat Ntn1 cKO mice (bottom panel). (G-H) mEPSC amplitude (G) and frequency (H) measured from fluorescently identified GABA neurons (n=30 controls, n=32 cKO, t=2.048, df=60, *p<0.05, and t=3.966, df=60, ***p<0.001). (I) Schematic of stimulating electrode placement in horizontal midbrain slice and example EPSCs. (J-K) Paired pulse ratio in dopamine (J, n=18 controls, n=21 cKO), or GABA neurons (K, n=14 controls, n=21 cKO).

Our observed reduction in mEPSC frequency suggests that loss of *Ntn1* function could act presynaptically, potentially through postsynaptic netrin-1 secretion^17^. To test potential presynaptic changes in vesicle release probability, we analyzed the paired-pulse ratio (PPR) of electrically evoked EPSCs delivered 50 ms apart. *Ntn1* mutagenesis in either dopamine or GABA neurons did not result in a significant change in PPR compared to controls, suggesting no measurable change in presynaptic release (Figure 3J-K).

Because netrin-1 is a secreted protein, it is also possible that *Ntn1* loss of function in one cell type could affect synaptic connectivity in adjacent neurons in which the gene was not inactivated, inducing a non-cell autonomous effect. To address this, we recorded mEPSCs from non-YFP-expressing (presumptively non-dopamine) neurons in DAT-Cre mice injected with *Ntn1* CRISPR or control virus, and from non-YFP-expressing (presumptively non-GABA) neurons in Vgat-Cre injected mice. We did not observe significant non-cell autonomous effects on mEPSCs from non-targeted cells (Figure 3 supplement 2). Similarly, we also did not observe non-cell autonomous effects on mIPSCs from non-targeted cells (Figure 3 supplement 2).

### *Ntn1* loss of function in VTA-dopamine neurons has little effect on behavior

Dopamine producing neurons of the VTA regulate multiple aspects of locomotor activity, motivated behavior, and psychomotor activation. To determine whether conditional mutagenesis of *Ntn1* in dopamine neurons, and subsequent reduction in excitatory synaptic connectivity impacts these behaviors, we injected DAT-Cre mice with AAV1-FLEX-SaCas9-sg*Ntn1* or AAV1-FLEX-SaCas9-sg*Rosa26* (control) and assayed them in multiple behavioral paradigms. First, we monitored day-night locomotion in control and AAV1-FLEX-SaCas9-sg*Ntn1* injected DAT-Cre mice. No significant differences were detected (Figure 4B and Figure 4 supplement 1).

**Figure 4.**
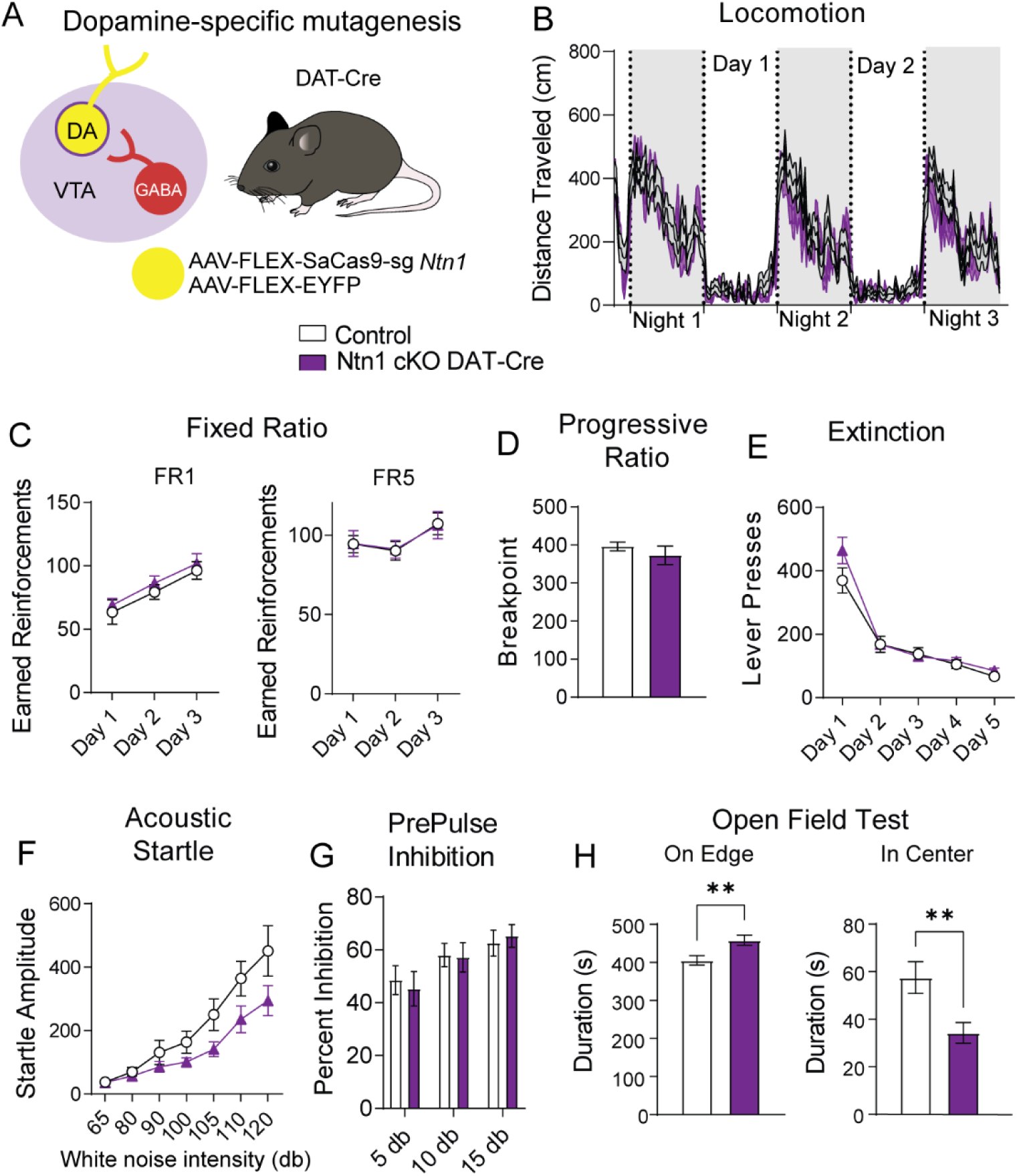
*Ntn1* cKO in DA neurons results in little behavioral alteration. (A) Schematic summarizing cell type specific knockout procedure. (B) Distance traveled in 15 min bins over the course of 3 nights and 2 days (n=21 control; n=15 cKO). (C) Earned reinforcers during 3 days of FR1 or FR5 operant conditioning. (D) Breakpoint (maximum presses per reinforcer) on a progressive ratio task. (E) Lever presses per session during 5 days of extinction training. (F) Acoustic startle response to varying intensity white noise stimuli (G) Percent inhibition of startle response following pre-pulse at indicated intensities. (H) Time on edge or in center of open field arena during a 10 minute test session (B-H: n=19 control; n=15 cKO. H Edge: t=2.897, df=32, **p<0.01, H Center: t=2.750, df=32, **p<0.01).

To determine whether appetitive conditioning behaviors are disrupted by loss of *Ntn1* function in VTA dopamine neurons, we assayed mice in a simple instrumental conditioning paradigm using a fixed-ratio 1 (FR1) followed by a fixed ratio 5 (FR5) schedule of reinforcement in which 1 or 5 lever presses are required to obtain a food reward, respectively. We did not observe significant differences in either of these behavioral tasks (Figure 4C). Next, we monitored motivated behavior using a progressive ratio schedule of reinforcement in which the number of lever presses required for reinforcement increases non-arithmetically (1, 2, 4, 7, 13, 19, 25, 34, 43, 52, 61, 73…), and again did not observe significant differences between control and experimental mice (Figure 4D). Following PR, we reinstated FR1 responding for 3 days followed by extinction training, and again did not detect any differences between the two groups (Figure 4E), indicating *Ntn1* loss of function in VTA dopamine neurons did not alter appetitive conditioning behaviors.

To determine whether sensory-motor gating is altered in mice with loss of *Ntn1* function in VTA dopamine neurons, we assayed them in acoustic startle and pre-pulse inhibition (PPI) paradigms. Although acoustic startle responses were reduced in AAV1-FLEX-SaCas9-sg*Ntn1* injected mice, this did not reach significance (Figure 4F). Moreover, we did not observe differences in PPI percentage inhibition (Figure 4G). These results indicate that loss of *Ntn1* function in VTA dopamine neurons does not appear to affect psychomotor activation.

In addition to reinforcement and motivation, dopamine regulates other dimensions of affective behavior. To test whether anxiety-related behavior is affected in experimental mice relative to control mice, we assayed them in an open-field test. AAV1-FLEX-SaCas9-sg*Ntn1* injected DAT-Cre mice spent significantly more time on the edge of the open field arena and significantly less time in the center of the arena, consistent with an elevation in anxiety-like behavior (Figure 4H).

### *Ntn1* loss of function in VTA-GABA neurons affects multiple behaviors

To determine whether reducing excitatory synaptic connectivity onto VTA GABA neurons through the loss of *Ntn1* function in these cells impacts behavior, we injected Vgat-Cre mice with AAV1-FLEX-SaCas9-sg*Ntn1* or AAV1-FLEX-SaCas9-sg*Rosa26* (control) into the VTA as described previously and tested these mice using the same behavioral paradigms described above. In contrast to *Ntn1* mutagenesis in dopamine neurons, this manipulation in VTA GABA neurons resulted in a significant increase in locomotor activity (Figure 5B and Figure 5 supplement 1).

**Figure 5:**
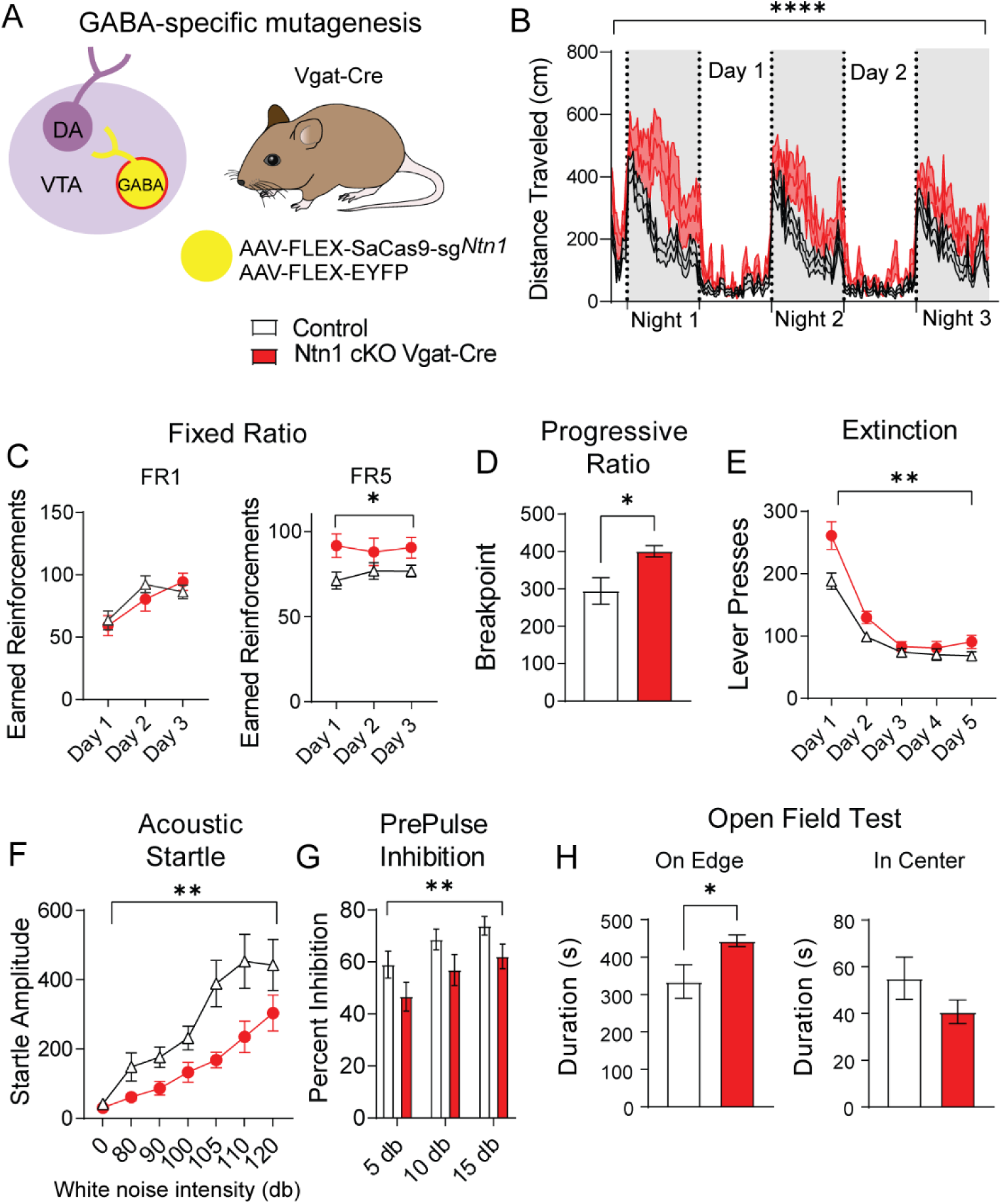
*Ntn1* cKO in GABA VTA neurons resulted in significant behavioral alterations. (A) Schematic summarizing cell type specific knockout procedure. (B) Distance traveled in 15 min bins over the course of 3 nights and 2 day (n=26 controls, n=23 cKO, Two-way ANOVA Group F (1, 11797) = 527.4, ****p<0.0001). (C) Earned reinforcers during 3 days of FR1 or FR5 operant conditioning (n=18 control; n=15 cKO; Two-way ANOVA F Group (1, 31) = 4.261, *p<0.05). (D) Breakpoint (maximum presses per reinforcer) on a progressive ratio task (t=2.577, df=31, *p<0.05). (E) Lever presses per session during 5 days of extinction training (Two-way ANOVA F Group F (1, 31) = 10.23, **p<0.01). (F) Acoustic startle response to varying intensity white noise stimuli (GABA-cKO; Group Factor F (1, 31) = 7.891, **p<0.0085, Startle x group F (6, 186) = 2.186, P=0.0462). (G) Percent inhibition of startle response following pre-pulse at indicated intensities (Group Factor F (1, 93) = 9.181, **p<0.01). (H) Time on edge or in center of open field arena during a 10 minute test session (edge: t=2.248, df=31, *p<0.05).

In the FR1 schedule of reinforcement we did not observe a significant difference between the groups; however, we observed an increase in the number of earned reinforcements in the FR5 schedule in mice with *Ntn1* loss of function in VTA GABA neurons (Figure 5C). We also observed an increase in the PR schedule of reinforcement in these mice relative to controls (Figure 5D). During extinction training, mice with *Ntn1* loss of function in VTA GABA neurons had a significant delay in the rate of extinction following reinstatement of FR1 conditioning (Figure 5E).

Analysis of sensory-motor gating in these mice revealed that Vgat-Cre mice injected with AAV1-FLEX-SaCas9-sgNtn1 had a significant reduction in the acoustic startle relative to control mice (Figure 5F) that was accompanied by a reduction in PPI (Figure 5G). Similar to mutagenesis of *Ntn1* in dopamine neurons, this manipulation in GABA neurons resulted in an increase in anxiety-like behavior as demonstrated by an increased time on edge; though we only observed a trend towards a reduction in time spent in the center of the open field arena (Figure 5H).

### Loss of netrin-1 in dopamine neurons largely reverses the effects of *Ntn1* mutagenesis in GABA neurons

A loss of netrin-1 in VTA-dopamine neurons resulted in decreased excitatory synaptic input to those cells (theoretically reducing dopamine activity) (Figure 6A), and loss of netrin-1 in VTA-GABA neurons resulted in decreased excitatory tone onto GABA neurons, which would be predicted to increase dopamine activity through disinhibition^18^ (Figure 6A). Based on these observations, we asked whether a loss *Ntn1* in both cell types would restore the balance of activity in the midbrain, or whether there is a hierarchical effect of *Ntn1* loss of function in GABA neurons. To address this, we crossed DAT-Cre with Vgat-Cre mice to develop a DAT-Cre::Vgat-Cre transgenic line, injected these mice with AAV1-FLEX-SaCas9-sg*Ntn1* or AAV1-FLEX-SaCas9-sg*Rosa26* (control)(Figure 6B), and assayed them using the previous behavioral battery.

**Figure 6:**
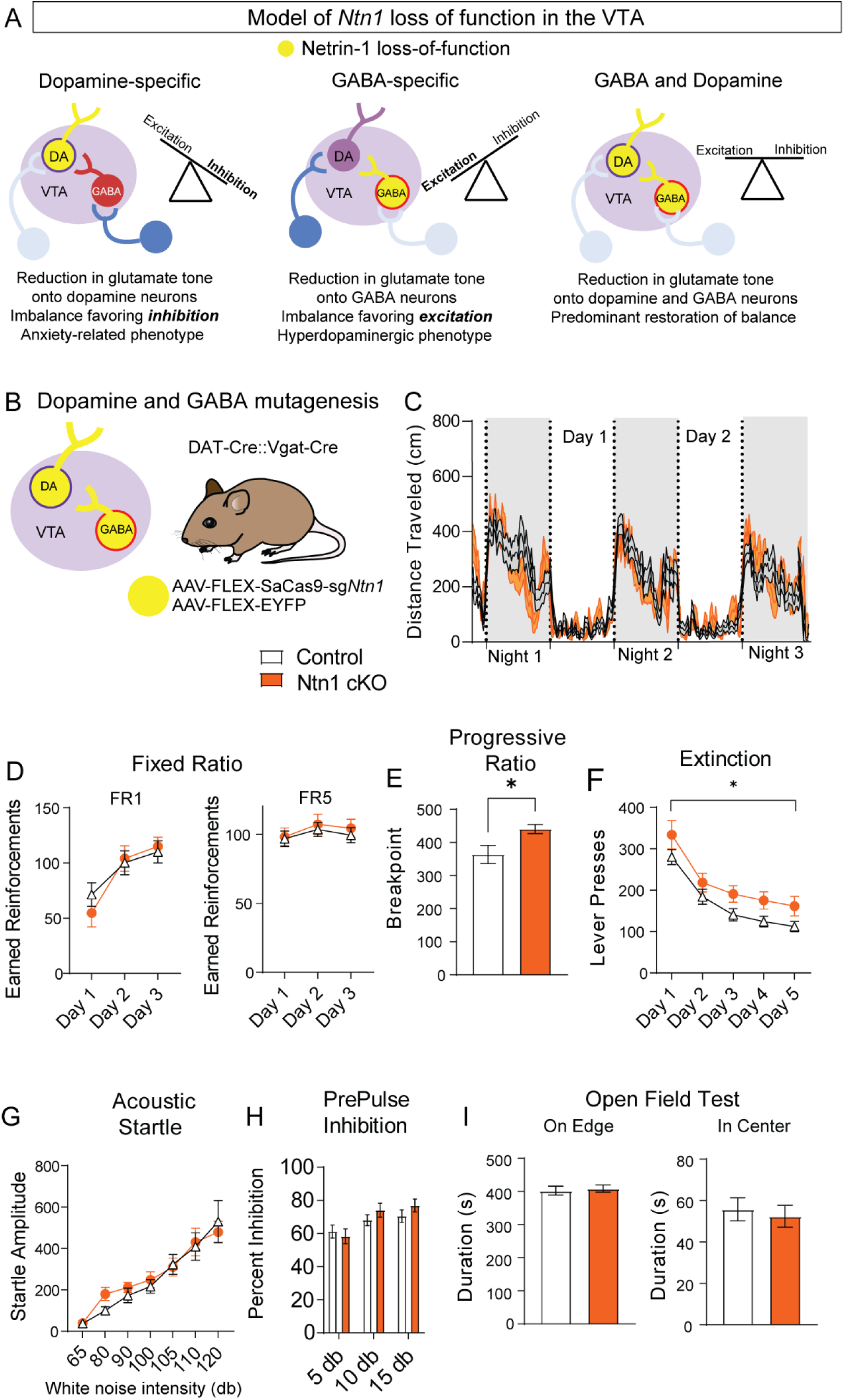
*Ntn1* cKO in DAT_IRES_::Vgat-Cre mice partially rescues behavioral phenotype. A) Model of *Ntn1* loss of function in the VTA on excitatory and inhibitory balance. B) Schematic of GABA and Dopamine Ntn1 cKO. C) Distance traveled in 15 min bins over the course of 3 nights and 2 day (Two-Way ANOVA Group Factor F (1, 45) = 0.01279, p<0.05). D) Earned reinforcers during 3 days of FR1 or FR5 operant conditioning. E) Breakpoint (maximum presses per reinforcer) on a progressive ratio task (n=21 controls, n=20 Ntn1 cKO, t=2.502, df=39, *p<0.05) F) Lever presses per session during 5 days of extinction training (Two-way ANOVA Group Factor F (1, 39) = 6.990, *p=0.0117). (G) Acoustic startle response to varying intensity white noise stimuli. G) Percent inhibition of startle response following pre-pulse at indicated intensities. (H) Time on edge or in center of open field arena during a 10 minute test session.

Simultaneous *Ntn1* loss of function in VTA GABA and dopamine neurons largely reversed the hyperlocomotor phenotype (Figure 6C) observed with *Ntn1* mutagenesis in VTA GABA neurons alone, though a modest, increase in daytime locomotion remained (Figure 6 supplemental figure 1). Similarly, loss of *Ntn1* in both VTA GABA and dopamine neurons resulted in operant responding during FR1 and FR5 that was similar to controls (Figure 6D). Motivation as measured in the PR task was elevated in the double transgenic Cre line following *Ntn1* mutagenesis (Figure 6E) and extinction was impaired (Figure 6F), though these phenotypes were less robust than those observed in the VTA GABA only mice. Finally, loss of netrin-1 in both cell types resulted in acoustic startle and PPI responses (Figure 6G-H), and open field activity (Figure 6I) that was similar to control mice.

## DISCUSION

Here we show that netrin-1 is present in both dopamine and GABA producing neurons of the adult VTA, and loss of netrin-1 function via genetic inactivation in either cell type results in a significant disruption of excitatory synaptic connectivity. The exact mechanisms by which netrin-1 regulates glutamatergic connectivity remains to be resolved. Likely mechanisms include netrin-1 regulation of the actin cytoskeleton and receptor transport vesicles through its activation of the cognate receptor DCC^19^. The latter is consistent with our observed decrease in the amplitude of mEPSCs and previous reports of netrin-1 regulating the delivery of GluA1 containing AMPA receptors to the postsynaptic density^17^. Our finding that mEPSC frequency, but not paired pulse ratio, were affected by netrin-1 loss further suggests netrin’s role in modulating excitatory synaptic connectivity is likely confined to postsynaptic mechanisms.

Loss of netrin-1 in dopamine neurons had little effect on behavior; however, we did observe and increase in anxiety-like behavior and measured by the open field assay consistent with the proposed role of dopamine in the modulation of anxiety-related behavior^20^. The general lack of effect of reduced glutamatergic synaptic connectivity on appetitive behavior, locomotion, and sensory-motor gating is consistent with previous observations that reduced glutamatergic signaling in dopamine neurons largely does not affect these behaviors^21,22^. In contrast, loss of netrin-1 in VTA GABA neurons had a significant effect on multiple behaviors including locomotion, motivation, and acoustic pre-pulse inhibition, all of which are consistent with a hyperdopaminergic phenotype and with previous reports that disrupting GABA neuron function in the VTA induces similar phenotypes^7,23^.

Given the robust nature of the behavioral effects observed following *Ntn1* mutagenesis in VTA GABA neurons, we were initially surprised that simultaneous loss of netrin-1 in both GABA and dopamine neurons largely rescued the observed hyperdopaminergic phenotype. These results suggest that a balance of glutamatergic signaling in these two cell types is essential for the normal functioning of the mesolimbic dopamine system (Figure 6A). This finding is similar to what has been reported previously in the striatum. Beutler et al, demonstrated that loss of NMDA receptor signaling in dopamine D1 receptor-expressing neurons prevented the development of amphetamine sensitization; however, inactivation of NMDA receptors in D1R and D2R-expressing medium spiny neurons reversed this phenotype^24^.

While our findings shed light on the role of netrin-1 in adult VTA neurons, the question remains as to which netrin-1 receptors may be involved. Indeed, though Dcc is considered to be its canonical receptor, netrin-1 is a known ligand for several additional receptors, including DSCAM, Neogenin, and Unc5 homologues A-D^19,25^. Previous work has identified the presence of both Dcc and Unc5c receptors in the adult VTA (often in the same cells)^8^. It is also interesting to note that during development of the spinal cord netrin-1 expression in the floor plate attracts commissural axons to the midline, but following the arrival of these axons at the floor plate, Unc5 expression increases to suppress the attractive actions of DCC signaling^26^.

Whether a similar relationship exists for the formation of nascent synapses and the maintenance of excitatory synapses occurs in the VTA will be important to resolve. Of further note, in addition to the role of netrin-1/Dcc/Unc5 signaling in the regulation of commissural axons crossing the midline, Slit/Robo signaling repels axons away from the floor plate^27,28^ setting up a push-pull relationship between these pathways. We previously demonstrated that Robo2 maintains inhibitory synaptic connectivity in the adult VTA^7^, suggesting the existence of another ‘push/pull’ relationship between these two pathways in which netrin/Dcc/Unc5 regulates excitation and Slit/Robo signaling regulates inhibition.

Mutations in *NTN1* (netrin-1) and *DCC* in humans and have been associated with several dopamine associated psychiatric conditions, including neurodevelopmental disorders such as schizophrenia^29–31^ and major depressive disorder^30,32–34^, as well as multiple neurodegenerative disorders^35–37^. Our findings that *Ntn1* plays a key role in maintaining excitatory connectivity in the adult midbrain and controlling the inhibitory/excitatory balance in this region highlights the importance of understanding these critical developmental signaling pathways in the adult nervous system that are likely important for therapeutic considerations in targeting these pathways.

## METHODS

### Mice

All procedures were approved and conduced in accordance with the guidelines of the University of Washington’s Institutional Animal Care and Use Committee. Mice were housed on a 12:12 light:dark cycle with *ad libitium* access to food and water, except when undergoing food restriction for operant behavioral conditioning. Approximately equal numbers of male and female mice were used. Mice were group housed (2-5 mice per cage). Mice injected with CRISPR/YFP were allowed 4-5 weeks recovery after surgery to allow for viral expression, mutagenesis, and protein turnover before any testing.

### Viruses

All adeno-associated viruses (AAV) were produced in house, as previously described^14^. CRISPR viruses employed for this research: AAV1-FLEX-SaCas9-U6-sgNtn1, AAV1-FLEX-SaCas9-U6-sgRosa26, AAV1-FLEX-YFP.

### Surgeries

All mice used were 8-10 weeks of age at time of surgery. Mice were inducted using isoflurane at 5.0% and held at 2% throughout the procedure. Mice were stereotaxically injected bilaterally into the VTA using the following coordinates in mm, relative to bregma: A/P: −3.25; M/L + 0.5; D/V: (−4.9) – (−4.4), total volume 0.5 µL into each side. A/P coordinates were adjusted for Bregma/Lambda distances using a correction factor of 4.2 mm.

### *In situ* hybridization

Male and female mice (n=2 each sex, 8-12 weeks old) were used to verify mRNA expression in the VTA using RNAscope (*2*). Brains were flash frozen in 2-methylbutane and representative coronal sections that spanned the VTA were sliced at 20 µm and slide mounted for hybridization. Sections were prepared for hybridization per manufacturer’s (Advanced Cell Diagnostics, Inc) instructions using probes for *Th* (Mm-*Th*), *Ntn1* (Mm-*Ntn1*-C2), and *Slc32a1* (Vgat; Mm-*Slc32a1*-C3). Slides were coverslipped with Fluoromount with DAPI (Souther Biotech) and imaged using a confocal fluorescent microscope (University of Washington Keck Center Leica SP8X confocal) and Keyence Fluorescence Microscope (Keyence). Quantification of colabeled cells was performed using CellProfiler, with thresholding and cell identification/overlap for each channel verified for each image manually prior to quantification.

### Immunohistochemistry

Mice were anesthetized with pentobarbitol and transcardially perfused with PBS followed by 4% PFA. Brains were post fixed for 24 hours in PFA at 4°C, followed by 48 hours in 30% sucrose. The VTA was coronally sectioned at 30 µm. Sections were kept in PBS with 0.3% Sodium Azide. Free floating sections were treated with 0.3% TBS-Triton-X 100 3×10 minutes, blocked in 3% Normal Donkey Serum for 1 hour, and treated overnight in primary antibody. Following 1-3 hours in secondary antibody (JacksonImmuno), sections were slide mounted and cover slipped with Fluoromount with DAPI. Images were collected on a Keyence Fluorescence Microscope (Keyence). For CRISPR validation, male and female DAT-Cre mice (8-12 weeks old) received AAV1-FLEX-SaCas9-U6-sgNtn1/ AAV1-FLEX-YFP (Ntn1-cKO) or AAV1-FLEX-SaCas9-U6-sgROSA26/AAV1-FLEX-YFP (controls) injections as described above, and quantification of colabeled cells for immunohistochemistry were performed using ImageJ 1.53 Cell Counter/Multi-point tool. Primary antibodies used: mouse anti-TH (1:1500, Millipore), chicken anti-Netrin-1 (1:1000, Abcam) and rabbit anti-HA (1:1500, Sigma)

### Slice electrophysiology

Mice injected with CRISPR/YFP were allowed 4-5 weeks recovery after surgery to allow for viral expression, mutagenesis and protein turnover. All solutions were continuously bubbled with O_2_/CO_2_. Horizontal (200 µm) brain slices were prepared from 12-20 week old mice in a slush NMDG cutting solution^38^ (in mM: 92 NMDG, 2.5 KCl, 1.25 NaH_2_PO_4_, 30 NaHCO_3_, 20 HEPES, 25 glucose, 2 thiourea, 5 Na-ascorbate, 3 Na-pyruvate, 0.5 CaCl_2_, 10 MgSO_4_, pH 7.3–7.4. Slices recovered for ∼12 min in the same solution warmed in 32°C water bath, then transferred to room temperature HEPES-aCSF solution (in mM: 92 NaCl, 2.5 KCl, 1.25 NaH_2_PO_4_, 30 NaHCO_3_, 20 HEPES, 25 glucose, 2 thiouria, 5 Na-ascorbate, 3 Na-pyruvate, 2 CaCl_2_, 2 MgSO_4_). Slices recovered for an additional 30 −60 min in HEPES solution at room temp. Whole-cell patch clamp recordings were made using an Axopatch 700B amplifier (Molecular Devices) using 3–5 MΩ electrodes. Recordings were made in aCSF (in mM: 126 NaCl, 2.5 KCl, 1.2 NaH_2_PO_4_, 1.2 MgCl_2,_ 11 D-glucose, 18 NaHCO_3_, 2.4 CaCl_2_) at 32°C continually perfused over slices at a rate of ∼1 ml/min. VTA dopamine and non-dopamine neurons were identified by fluorescence.

#### mE/IPSC

For miniature excitatory postsynaptic currents (mEPSCs), internal solution contained: 130 mM K-gluconate, 10 mM HEPES, 5 mM NaCl, 1 mM EGTA, 5 mM Mg-ATP, 0.5 mM Na-GTP. Picrotoxin (200 μM) was added to ACSF to block GABA_A_ receptor-mediated events. For miniature inhibitory postsynaptic currents (mIPSCs), internal solution contained: 135 mM KCl, 12 mM NaCl, 0.05 mM EGTA, 100 mM HEPES, 0.2 mM Mg-ATP, 0.02, and Na-GTP mM. To block glutamatergic events, 2 mM kynurenic acid was bath applied in the ACSF. All mIPSCs and mEPSCs cells were recorded in the presence of 1 mM tetrodoxin (TTX) to block action potentials. Cells were held at −60 mV for a minimum of 5 minutes prior to data acquisition. Data were analyzed using Clampfit 10.3 (pCLAMP 11 Software Suite, Molecular Instruments).

#### Paired Pulse Ratio

For PPR, internal solution contained: 130 mM K-gluconate, 10 mM HEPES, 5 mM NaCl, 1 mM EGTA, 5 mM Mg-ATP, 0.5 mM Na-GTP. Picrotoxin (200 μM) was added to ACSF to block GABA_A_ receptor-mediated events. Electrical stimulation was delivered using a concentric bipolar electrode placed rostral to the VTA. Data were analyzed using Clampfit 10.3 (pCLAMP 11 Software Suite, Molecular Instruments).

### Behavior

#### Locomotor activity

4 weeks after surgery, baseline locomotion was measured using locomotion chambers (Columbus instruments) that use infrared beam breaks to calculate ambulatory activity. Mice were singly housed in Allentown cages with reduced corncob bedding and provided with *ad libitum* access to food and water. Locomotion was monitored continuously for 3 nights 2 days

#### Open field testing

Mice were placed in a large circular arena (120 cm diameter) and activity was recorded for a period of 10 min using Ethovision software. Time in center, time on edge, and total distance were calculated.

#### Operant conditioning

Mice were tested on an operant conditioning paradigm in Med Associates boxes in the following order: FR1, FR5, Progressive Ratio, Reinstatement and Extinction. Each fixed ratio 1 (FR1) session lasted for 60 min. Levers were extended and remained extended until a lever press. Upon a lever press, levers were retracted and a sucrose pellet was immediately delivered into the food hopper. The levers did not extend again until the mouse made a head entry into the food hopper to retrieve the pellet. Reinforced FR1 sessions lasted for 3 days, followed by 3 days of FR5 (5 lever presses required to obtain sucrose pellet), and a single day of progressive ratio where the number of lever presses necessary for sucrose pellet delivery increases non-arithmetically (i.e., 1, 2, 4, 6, 9, 13…) over the course of the session. The progressive ratio session ended after 3 consecutive min of no lever presses or after 3 hours. After progressive ratio, mice again underwent FR1 reinforced training, followed by extinction for 60 min each session for five days. Here, levers extend and retract similarly to the FR1 reinforced paradigm, yet a sucrose pellet reward is omitted.

#### Acoustic startle and Prepulse inhibition

Acoustic startle responses were measured using acoustic startle chambers (San Diego Instruments). Prior to testing mice received a 10-min habituation period. Background noise was maintained at 65 dB throughout testing. After habituation, mice were presented with 5, 40-ms duration 120 dB, pulse-alone trials to obtain baseline startle responses, followed by 50 trials of either a startle pulse-alone, 1 of 3 prepulse trials, or a null trial, in which no acoustic stimulus is presented. Startle trials consisted of a 40 ms, 120-dB pulse of white noise. The 3 prepulse trials consisted of a 20-ms prepulse of 70-, 75-, or 80-dB intensity (5, 10, and 15 dB above background) that preceded 120-dB startle pulse by 100 ms. Peak amplitude of the startle response (65 ms after pulse onset) was used as the measure of startle response magnitude.

## Statistics

Data were analyzed for statistical significance using GraphPad Prism. All statistical tests were two-sided and corrected for multiple comparisons where appropriate.

## Acknowledgements

We would like to thank the staff of the University of Washington’s Comparative Medicine Animal Facilities, the University of Washington’s Keck Imaging Center, and the administrative staff of the Molecular and Cellular Biology Graduate Program.

## Author contributions

MMC, MS and LSZ conceived and designed experiments. MMC, ACH, and LZ designed and generated AAVs. MMC and BJ performed behavioral testing and analysis. MMC performed all surgeries and in situ hybridization experiments. MMC and MS performed and analyzed electrophysiology experiments. GE and BH performed and analyzed immunohistochemistry and viral effectiveness images. All authors provided input and approval of the manuscript.

## Supplemental Figures

**Figure 3 supplemental 1:**
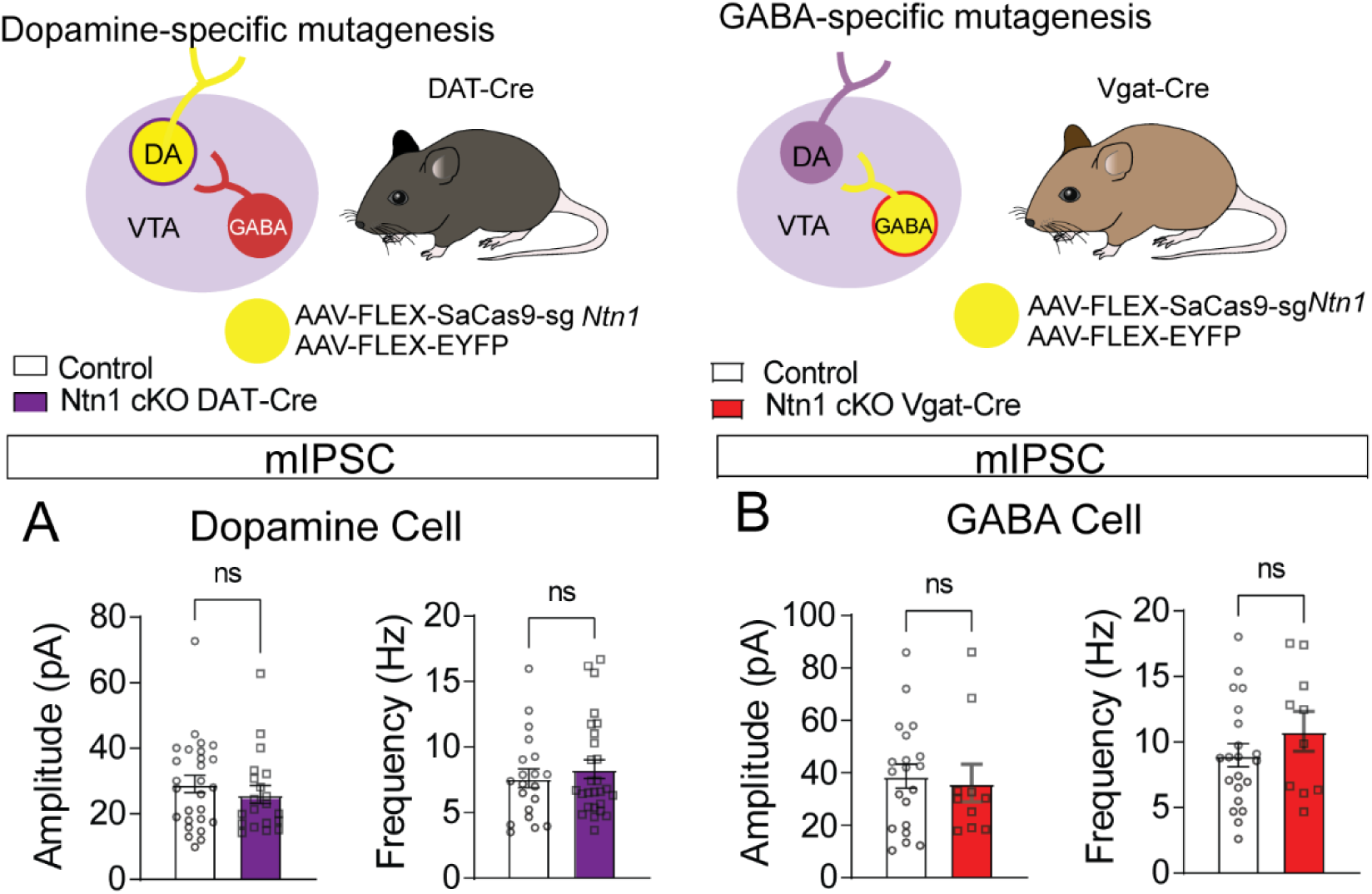
No significant differences in inhibitory synaptic connectivity associated with *Ntn1* loss of function. A) mIPSC amplitude and frequency of DAT-Cre fluorescently identified dopamine neurons (n=20 controls, n=27 cKO). B) mIPSC amplitude and frequency of Vgat-Cre fluorescently identified GABA neurons (n=20 controls, n=10 cKO).

**Figure 3 supplement 2:**
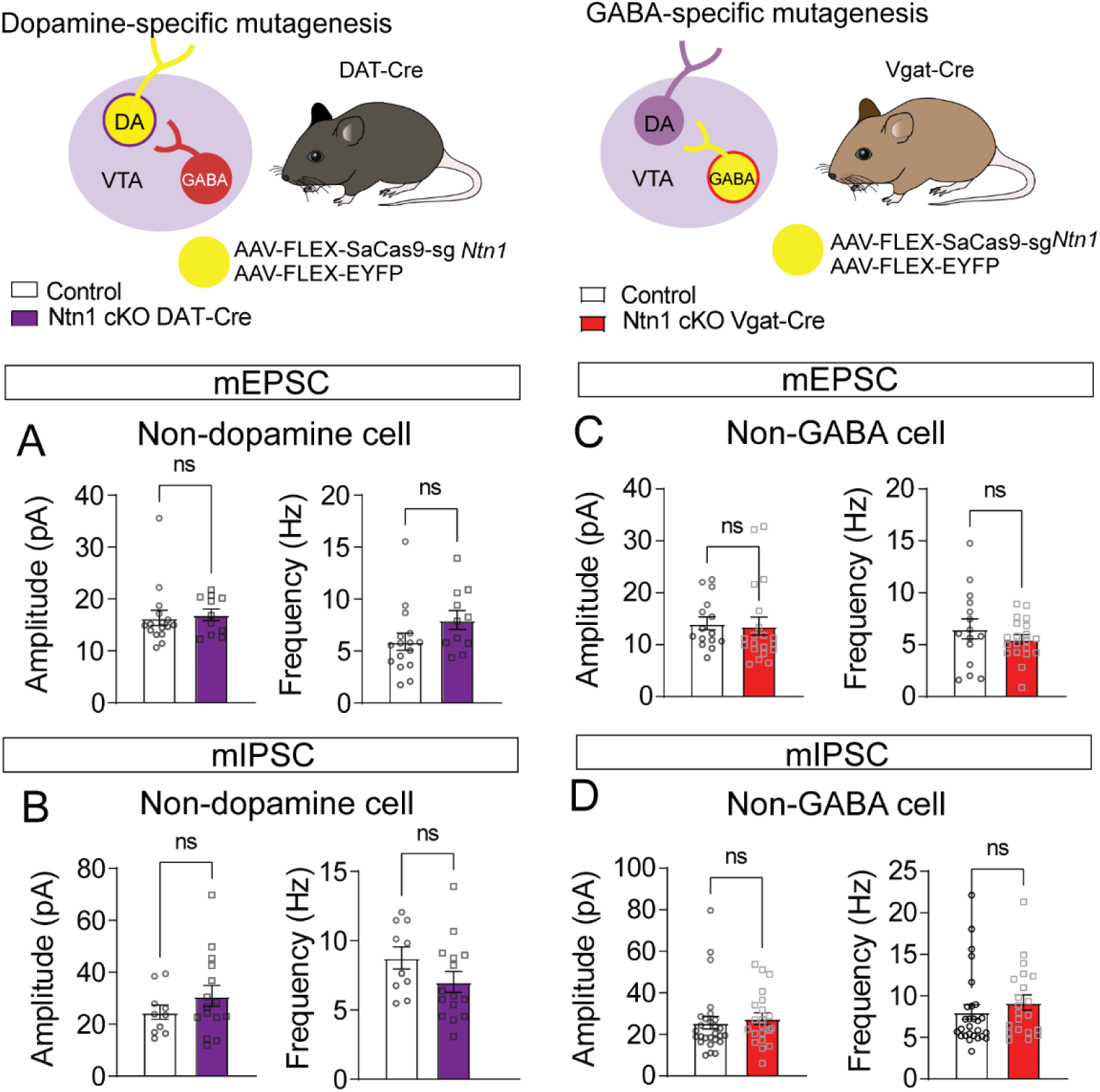
No significant differences in excitatory or inhibitory synaptic connectivity in non-targeted cell types. A) mEPSC amplitude and frequency recorded from non-fluorescent cells in DAT-Cre mice (presumptively non-dopamine neurons) (n=16 controls, n=11 cKO). B) mIPSC amplitude and frequency recorded from non-fluorescent cells in DAT-Cre mice (n=10 controls, n=15 cKO). C) mEPSC amplitude and frequency recorded from non-fluorescent cells in Vgat-Cre mice (presumptively non-GABA neurons) (n=18 controls, n=20 cKO). D) mIPSC amplitude and frequency recorded non-fluorescent cells in Vgat-Cre mice (n=27 controls, n=21 cKO).

**Figure 4 supplement 1:**
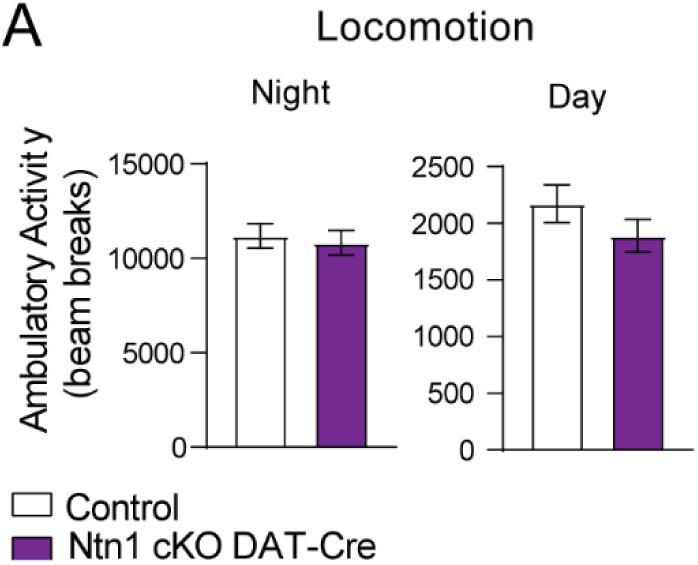
Average day and night locomotion in DAT-Cre mice. A) Ambulatory activity (beam breaks) averaged across 3 nights and 2 days (n=21 control; n=18 cKO).

**Figure 5 supplement 1:**
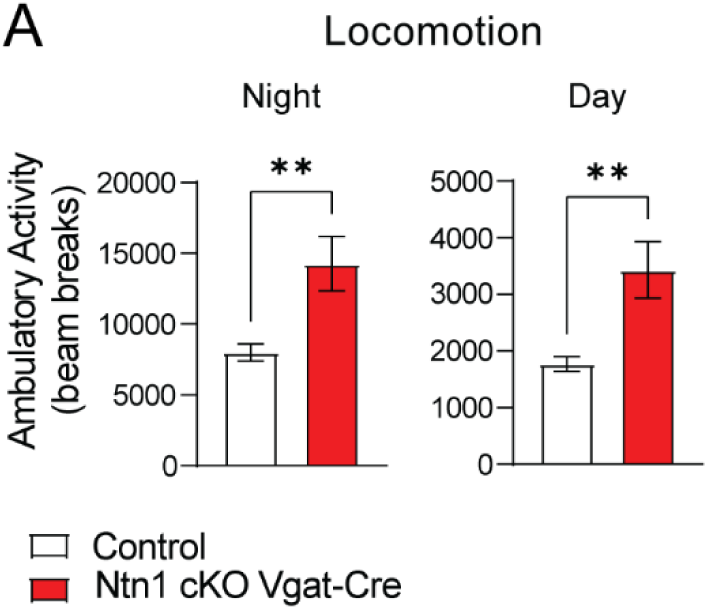
Average day and night locomotion in Vgat-Cre mice. A) Ambulatory activity (beam breaks) averaged across 3 nights and 2 days (n=26 control; n=26 cKO, (t=3.109, df=50 **p=0.01 and t=3.227, df=50 **p=0.01).

**Figure 6 supplement 1:**
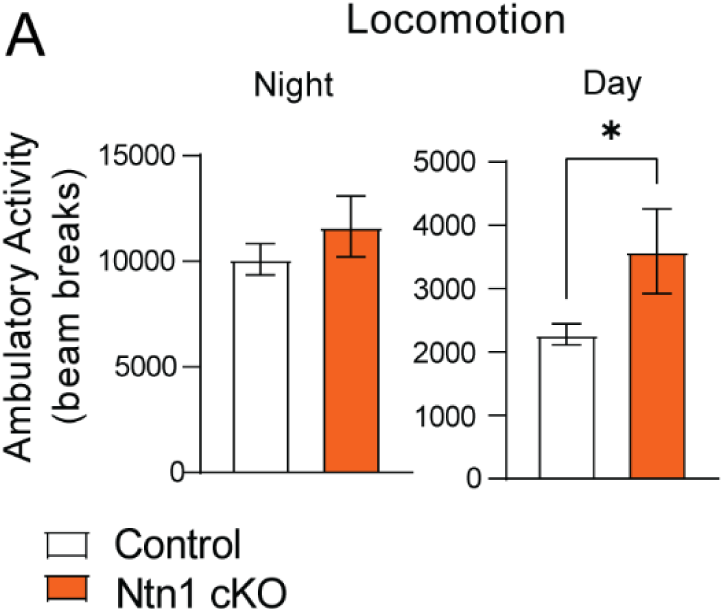
Average day and night locomotion in DAT-Cre::Vgat-Cre mice. A) Ambulatory activity (beam breaks) averaged across 3 nights and 2 days (n=26 controls n=21 cKO, t=2.091, df=45 *p<0.05).

